# How the brain represents a romantic partner: dissociable roles of the nucleus accumbens and anterior insula

**DOI:** 10.64898/2026.02.16.706085

**Authors:** Kenji Fujisaki, Ryuhei Ueda, Ryusuke Nakai, Nobuhito Abe

## Abstract

Humans form selective and enduring pair bonds with romantic partners, a principal feature of human sociality. Neuroimaging studies have shown that romantic partners are differentially represented from other individuals in the nucleus accumbens (NAcc) and anterior insula (aINS), and that the specificity of partner representations in the NAcc diminishes as relationships mature. However, it remains unclear whether such differentiation reflects partner-specific coding or mere differences in familiarity with others, and whether these regions play different roles in romantic bonding. To address these questions, we applied multiple regression representational similarity analysis to fMRI data from 51 heterosexual male participants in early romantic relationships. The data were acquired during a social incentive delay task, in which participants anticipated social approval from their female romantic partner, a female friend, or an unfamiliar female individual. This approach allowed us to dissociate partner-specific representations from familiarity-related effects in the NAcc and aINS. We found that both regions exhibited partner-specific representations that could not be explained by familiarity. Consistent with previous findings, partner specificity in the NAcc was negatively associated with relationship duration, indicating that partner-specific coding in this region is established early in romantic relationships and diminishes as relationships progress. Moreover, greater partner specificity in the aINS was associated with more frequent intrusive thoughts about the partner. Together, these findings demonstrate that romantic partners are represented in the NAcc and aINS in a qualitatively distinct manner from other individuals, and that these regions support dissociable aspects of romantic bonding.

**Key Points:**

- Multiple regression representational similarity analysis revealed partner-specific representations in the nucleus accumbens and anterior insula that cannot be explained by familiarity.
- Individuals in longer relationships showed reduced partner specificity in the nucleus accumbens, consistent with prior findings.
- Individuals exhibiting greater partner specificity in the anterior insula reported more frequent intrusive thoughts about their partner, indicating dissociable psychological functions across regions.

## Introduction

Humans form selective, enduring pair bonds with romantic partners—a trait rare among mammals (Lukas & Clutton-Brock, 2013) that has been argued to contribute to the evolution of distinctively human life history and social organization (Coxworth et al., 2015; Gavrilets, 2012; Shultz et al., 2011). Unlike most mammals, where proximity between adult heterosexual pairs is transient and tied to reproductive timing (Clutton-Brock, 1989; Kleiman, 1977), humans’ committed relationship with specific romantic partners can last for decades. Understanding the neural mechanisms that distinguish and selectively prioritize a romantic partner over other individuals is therefore fundamental to explaining a core feature of human sociality. Moreover, because romantic relationships profoundly affect mental and physical health (Hudson et al., 2020; Lawrence et al., 2019), identifying the neural basis of partner bonding has societal and clinical relevance.

Building on this background, studies have examined the neural basis of romantic love using neuroimaging techniques, particularly functional magnetic resonance imaging (fMRI). Early fMRI studies have demonstrated greater activity toward partners in reward-and motivation-related regions, such as the ventral tegmental area, caudate nucleus, and nucleus accumbens (NAcc), as well as in the saliency network, including the anterior cingulate cortex and insula (Acevedo et al., 2012; Aron et al., 2005; Bartels & Zeki, 2000; Fisher et al., 2010; Rafi et al., 2020; Xu X. et al., 2011). Extending this body of work, our recent studies examined partner-related processing at a finer representational level using multivoxel pattern analysis (Cohen et al., 2017). Our first study showed that neural representations of romantic partners could be reliably decoded from those of an unfamiliar attractive individual by analyzing the voxelwise brain activity patterns within the NAcc (Ueda & Abe, 2021). In subsequent work, we found that the NAcc and anterior insula (aINS) differentiated female partners from female friends, and that female friends were represented more similarly to male friends than to partners (Fujisaki et al., 2026). Notably, we also observed that such partner specificity in the NAcc was negatively associated with relationship length, suggesting dynamic changes in partner-related processing as relationships mature.

Despite these advances, two critical questions remain unresolved. First, our prior studies (Fujisaki et al., 2026; Ueda & Abe, 2021) did not explicitly control for familiarity, making it unclear whether the observed neural distinctions reflect partner-specific coding or simply greater familiarity with romantic partners relative to comparison targets (i.e., unfamiliar individuals or friends). Resolving this ambiguity is theoretically critical: partner-specific coding would indicate specialized mechanisms for romantic bonding, whereas familiarity-based coding would suggest that romantic relationships recruit the same neural architecture supporting other close bonds. A recent meta-analysis has shown that both the NAcc and aINS respond preferentially to familiar and socially close individuals across relationship types, including romantic partners, close friends, and family members (Bortolini et al., 2024). Considering these findings, explicit experimental control is necessary to address the present question.

Second, although Fujisaki et al. (2026) have observed distinct partner representations in the NAcc and aINS, further investigations are required to elucidate their functional roles in romantic bonding. Based on the role of NAcc in assigning incentive value to stimuli that draws attention and promotes approach behavior (Berridge & Robinson, 1998; Salamone & Correa, 2012), and evidence from prairie voles showing that this region causally supports partner preference and pair bond formation (Feldman, 2017; Walum & Young, 2018), partner representations observed in the NAcc likely reflect selective preference and approach motivation toward one’s partner.

In contrast, the aINS might support intrusive and obsessive thoughts about the romantic partner, which are typically observed in the early stage of romantic relationships (Langeslag et al., 2012; Nilakantan et al., 2014). This prediction is informed by evidence that the aINS plays a role in integration of internal and external signals to represent subjective salience, as well as in recalling and holding such representations in mind (Craig, 2009; Menon & Uddin, 2010; Naqvi et al., 2014; Naqvi & Bechara, 2010). Converging evidence from substance addiction further indicates that the aINS is critical for the conscious experience of urges and cravings (Naqvi et al., 2007; Naqvi & Bechara, 2009), suggesting a role in maintaining mental focus on salient targets. Furthermore, persistent and involuntary cognition centered on a specific romantic partner bears marked phenomenological similarity to obsessive–compulsive disorder (OCD) (Doron et al., 2014; Leckman et al., 1999; Thompson et al., 2020). Importantly, neuroimaging studies have consistently implicated the aINS as part of the neural circuitry underlying OCD (de Wit et al., 2014; Norman et al., 2019; Yu et al., 2022). Together, these observations raise the possibility that the aINS contributes to romantic bonding by maintaining partner-related representations with heightened salience, thereby promoting the spontaneous intrusion of partner-related thoughts.

Here, we address these two issues. First, to determine whether the NAcc and aINS surely carry specific representations of the romantic partner rather than familiarity, we applied multiple regression representational similarity analysis (RSA; Chikazoe et al., 2014) to spatial patterns of neural activity in response to female romantic partners, female friends, and unfamiliar female individuals in male participants. This approach allowed us to statistically control for familiarity and isolate the independent contribution of partner-specific representations in both the NAcc and the aINS. Furthermore, to clarify the functional roles of these two regions, we conducted correlation analyses to test two hypotheses. First, we examined whether partner specificity in the NAcc varies with relationship duration at relatively early stages of romantic relationships, which was observed in previous work (Fujisaki et al., 2026). Second, we tested whether individual differences in the frequency of intrusive thoughts about one’s partner are associated with the distinctiveness of partner-representations in the aINS.

## Methods

### Participants

We initially enrolled 55 male participants to meet our a priori target sample size. An a priori power analysis was conducted using G*Power (Version 3.1.9.6; Faul et al., 2007) to determine the required sample size for testing whether the two regression coefficients of interest estimated from the multiple regression RSA were greater than zero. This analysis was based on a one-tailed one-sample *t* test with a Bonferroni-corrected α = 0.025 (0.05/2), 90% power, and an assumed medium effect size (Cohen’s *d* = 0.5), indicating that a minimum of 44 participants was required. To account for potential attrition of approximately 20%, based on our prior work (Ueda & Abe, 2021) and the longitudinal nature of the present study involving a 10-day experience-sampling procedure, we decided to recruit 55 participants.

Participants were recruited to meet the following inclusion criteria: (a) male sex assigned at birth, (b) age between 20 and 29 years, (c) in a romantic relationship with a female partner for no more than six months at the time of the experiment, (d) unmarried and without children, (e) right-handed, (f) normal or corrected-to-normal vision, and (g) no history of neurological or psychiatric disorders. We restricted the sample to males aged 20–29 to minimize potential confounding effects of sex differences and age-related variability, and to maintain consistency with previous studies (Fujisaki et al., 2026; Ueda & Abe, 2021). Relationship duration was limited to within six months to focus on the early stage of romantic love, which has been characterized by frequent intrusive and obsessive partner-related thoughts during the first several months of a relationship (Langeslag et al., 2012; Marazziti et al., 1999; Nilakantan et al., 2014). One participant exceeded the relationship-length criterion by one month (7 months) at the time of participation but was still retained in the sample. All participants provided written informed consent after receiving a full explanation of the study, in accordance with the guidelines approved by the Kyoto University Ethics Committee.

Four participants were excluded from the final analysis. One participant could not undergo scanning because of a fixed orthodontic bridge incompatible with MRI safety standards, two did not identify as heterosexual in post-experiment questionnaires, and one exceeded the mean + 3 SD cutoff for error trials in the social incentive delay task. The final sample comprised 51 participants (mean age = 21.4 years, SD = 1.6, median = 21, range = 20–28; mean relationship length = 4.4 months, SD = 1.5, median = 5, range = 1–7).

### Stimuli

Following the procedure of our previous study (Ueda & Abe, 2021), we prepared video clips and facial images depicting three targets, all of whom were assigned female at birth: the participant’s romantic partner, a female friend of the participant, and an attractive female individual unfamiliar to the participant. Stimuli for the romantic partner and the female friend were unique to each participant, as they depicted each participant’s own partner and friend, whereas stimuli for the unfamiliar female were obtained from a single highly attractive individual whose stimuli had been created and validated in our previous study (Ueda & Abe, 2021) and were identical across participants. These stimuli were used in both the social incentive delay task and the subsequent rating task. To minimize variability in the nature of friendships, participants were instructed to nominate a female friend aged 20–29 years for whom they did not experience romantic interest. Both romantic partners and friends provided written informed consent and took part in the recording sessions without the presence of the fMRI participant.

Video clips conveying social approval were recorded for each individual and served as positive feedback following successful responses in the social incentive delay task. Specifically, we created ten short clips per individual, in which individuals displayed smiling facial expressions accompanied by positive hand gestures (e.g., making a V sign, an OK sign, or waving their right hand toward the camera). To ensure consistency across stimuli, individuals were instructed to make facial expressions and gestures imitating sample images. For nonapproval feedback presented following error responses in the social incentive delay task, we created a video clip of each person displaying a neutral facial expression without any gestures. From each of these clips, a still image of the neutral face was extracted and used as the cue stimulus in the task. All video clips and images were edited using Adobe Premiere Pro and Photoshop.

### Social Incentive Delay Task

Participants completed a modified version of the social incentive delay task during fMRI scanning (Ueda & Abe, 2021), which was originally proposed by Spreckelmeyer et al. (2009). The social incentive delay task is a modified version of the monetary incentive delay task (Knutson et al., 2000) and widely used to investigate the neural mechanisms underlying social reward processing (Cromwell et al., 2020). In this task, participants are required to respond quickly to a target stimulus to receive positive feedback indicating social approval. A brief anticipatory delay without any stimuli depicting the target person precedes the feedback, reducing the potential influence of facial recognition processes and minimizing confounding effects related to deliberate evaluation. The task was programmed and presented using the Presentation software package (Version 23.0; Neurobehavioral Systems, Inc., Berkeley, CA, www.neurobs.com). A schematic illustration of the task is shown in Figure 1.

**Figure 1.**
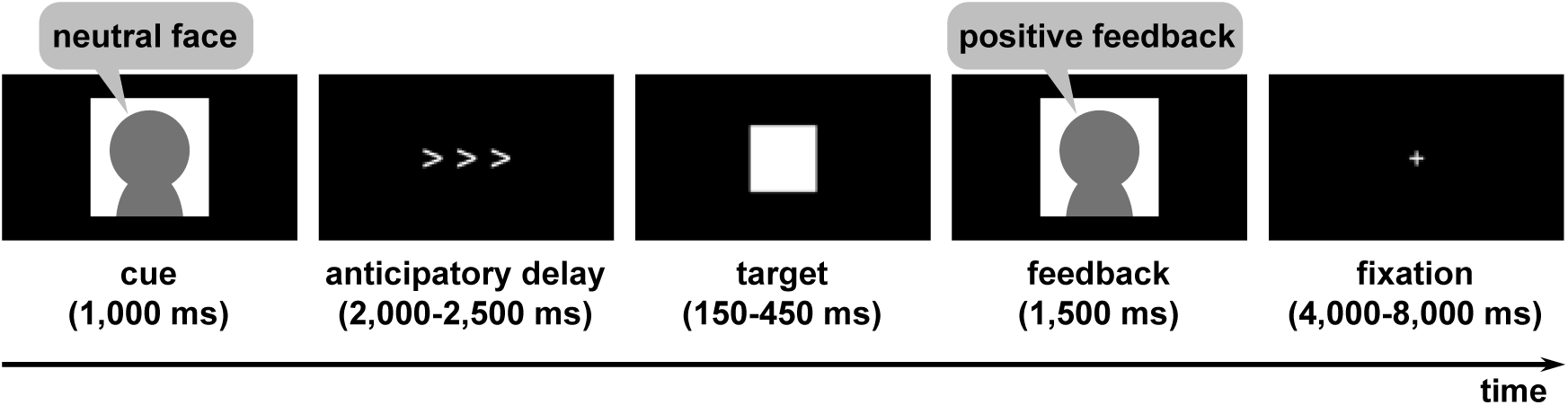
Schematic illustration of the social incentive delay task. In each trial, participants were first presented with a cue stimulus indicating the social target (romantic partner, female friend, or unfamiliar female individual). This cue was followed by an anticipatory delay phase with a variable duration. Subsequently, a white square target appeared briefly, during which participants were required to press a button. Successful responses were followed by a short video in which the cued person displayed a smile and a positive gesture, whereas unsuccessful responses were followed by a video showing a neutral facial expression.

In this study, the social incentive delay task had three conditions: the participant’s romantic partner (*partner* condition), a female friend of the participant (*friend* condition), and an attractive female individual unfamiliar to the participant (*unfamiliar* condition). The task comprised six runs of 30 trials (three conditions × 10 trials) in a pseudorandomized order, ensuring that no condition appeared more than twice in a row. Each trial started with a 1,000 ms cue, followed by a visual object indicating that the target stimulus would appear after a variable anticipatory delay (2,000–2,500 ms). A white square target was then presented for a variable duration (150–450 ms). Participants responded by pressing a button with their right index finger as quickly as possible when the target stimulus appeared. In the subsequent outcome phase, a hit (i.e., responding during target presentation) resulted in the presentation of a short video clip (1,500 ms) showing the cued person smiling and making a positive nonverbal gesture (see Stimuli). To enhance engagement, each run included 10 different video clips per person, each shown only once in a pseudorandom order. In contrast, a miss (i.e., responding after the target disappeared) or an error (i.e., responding before the target appeared or not responding at all) resulted in the presentation of a single video (1,500 ms) showing the cued person with a neutral facial expression and no nonverbal gestures. Unlike hit trials, only one neutral video per condition was used repeatedly for these outcomes. Each trial ended with a fixation cross presented for a variable intertrial interval (4,000–8,000 ms). To maintain comparable task difficulty across participants, an adaptive algorithm adjusted the target duration in 25-ms increments: shortening it when the hit rate exceeded 66% and lengthening it when the rate fell below 66% (Fujisaki et al., 2026; Ueda & Abe, 2021). For the first trial of the first run, the duration was fixed at 300 ms. In subsequent runs, the duration for the first trial carried over from the last trial of the previous run. The mean hit rate across participants was 61.6% (SD = 2.7%), and the mean number of error trials was 5.0 (SD = 5.9). No participant scored below the mean − 3 SD cutoff for hit rate (53.5%). However, one participant exceeded the mean + 3 SD cutoff (22.6) for error trials and was excluded from all analyses.

### Postscan Rating Tasks and Questionnaires

After the social incentive delay task, participants performed rating tasks outside the scanner. The task was implemented using PsychoPy Version 2022.2.4 (Peirce et al., 2019). They first rated the likeability of 30 video clips (10 per condition), featuring their own partner, a friend, and the unfamiliar female, which had been used as positive feedback stimuli in the social incentive delay task. In a pseudorandomized order, videos were shown for 1,500 ms, followed by a 1,000-ms fixation cross. Next, they rated their romantic interest in the same three individuals based on facial images with neutral expressions. All ratings were made on visual analog scales ranging from 0 (not at all) to 100 (extremely). Likeability ratings were *z*-scored within each participant across the 30 video clips, and mean *z*-scores for each condition (partner, friend, unfamiliar) were used for group-level analysis. We also asked whether participants personally knew the individual presented in the unfamiliar condition and confirmed that none reported knowing her.

Following the rating tasks, the participants completed a set of questionnaires. These included demographic information (age and sex), the Japanese version of the Triangular Love Scale (Kanemasa & Daibo, 2003; Sternberg, 1986), questions about whether they had an argument with their partner in the week before the experiment and whether any important changes had occurred in their relationship in the week before the experiment (Scheele et al., 2013), relationship length with the current partner, cohabitation status, the number of days since they last saw their partner, relationship history (operationalized as the number of prior heterosexual romantic relationships and, if more than one, the start and end dates of the most recent prior relationship), and sexual orientation. The Triangular Love Scale was used to assess romantic love toward the partner, conceptualized as comprising three components: intimacy, passion, and commitment.

### Assessment of intrusive thoughts

Starting on the day following the fMRI experiment, participants were asked to complete a follow-up survey over a 10-day period to assess intrusive thoughts about their partner. This assessment was conducted using an experience sampling method implemented via *exkuma*, a smartphone-based platform for experience sampling research (https://exkuma.com/). Participants received seven prompts per day delivered through their SMS application at pseudorandom times between 8:00 and 23:00, with a minimum interval of 1 hour between prompts, resulting in 70 assessments. At each prompt, participants reported the content of the thoughts they had just before accessing the prompt by selecting all applicable categories from a predefined list, including thoughts about their romantic partner, friends, family members, work or study, and other topics. Instead of directly asking whether participants had been thinking about their partner, responses were obtained through a forced-choice format, which minimized potential response bias. Participants also indicated whether they were currently spending time with someone (e.g., partner, friend, family member, or no one). An intrusive thought score was computed as the proportion of responses to prompts in which participants reported thinking about their romantic partner, excluding missing responses. Higher scores therefore reflected a greater frequency of partner-related thoughts during daily life.

### Behavioral Data Analysis

The following behavioral measures were statistically analyzed: response times during the social incentive delay task, and post-scan ratings of likeability and romantic interest. For these measures, we conducted a one-way repeated-measures analysis of variance with condition (partner, friend, unfamiliar) as a within-subject factor. When a significant main effect of condition was observed, we performed post hoc paired *t* tests with Bonferroni correction (adjusted *p* = *p* × 3). For romantic interest ratings, one participant did not provide complete data, and analyses were therefore conducted on fifty participants.

Throughout the study, statistical analyses were conducted using R (Version 4.5.0; R Core Team, 2025). Effect sizes were reported as partial *η*² for analyses of variance and as Hedges’ *g* for *t* tests, implemented in R using the *effectsize* package (Version 1.0.1; Ben-Shachar et al., 2020).

### fMRI Data Acquisition

Scanning was performed using a 3.0 T MAGNETOM Verio MRI scanner (Siemens Healthineers, Erlangen, Germany) with a 32-channel head coil. Participants lay in a supine position, with the head stabilized using foam padding to minimize motion. MRI-compatible glasses were provided to participants with inadequate unaided vision to ensure optimal viewing of visual stimuli. Stimuli were projected onto a screen and presented via a mirror mounted on the head coil. Button responses were recorded using a fiber-optic response box. Functional imaging was conducted using a T2*-weighted echo‒planar imaging (EPI) sequence sensitive to blood-oxygen-level-dependent (BOLD) contrast, with accelerated multiband acquisition to improve temporal resolution (Feinberg et al., 2010; Moeller et al., 2010; J. Xu et al., 2013). Functional slices were aligned to the anterior commissure–posterior commissure (AC–PC) line to standardize anatomical orientation across participants. The functional images were obtained with the following parameters: repetition time (TR) = 2,000 ms, echo time (TE) = 43 ms, flip angle = 80°, acquisition matrix = 96 × 96, field of view (FOV) = 192 × 192 mm, in-plane resolution = 2 × 2 mm, number of slices = 76, slice thickness = 2.0 mm with no interslice gap (interleaved acquisition), and multiband acceleration factor = 4. The first five volumes of the 173 samples acquired per run were discarded from analysis to allow for magnetic field stabilization. A high-resolution structural image was also acquired using a T1-weighted magnetization-prepared rapid gradient-echo (MPRAGE) pulse sequence, with a spatial resolution of 1 × 1 × 1 mm.

### fMRI Data Preprocessing and General Linear Model (GLM)

We conducted image preprocessing and statistical analyses using the Statistical Parametric Mapping software package (SPM 25; Wellcome Department of Cognitive Neurology, Institute of Neurology, London, England). Functional images were first slice-time corrected using the acquisition timing of the middle slice (952.5 ms) as a reference and subsequently realigned to correct for head motion across scans. Each participant’s T1-weighted structural image was coregistered to the mean functional image obtained during realignment and spatially normalized to the Montreal Neurological Institute (MNI) standard space using unified segmentation (Ashburner & Friston, 2005). The realigned functional images were then normalized using the same transformation parameters and resampled to 2 mm isotropic voxels. We used the resulting normalized, unsmoothed functional images for statistical analysis.

We designed a general linear model (GLM) for each participant to model the BOLD signal of each voxel, with regressors created separately for each run. For each run, the GLM contained three regressors representing the anticipatory delay phases for the three conditions (partner, friend, and unfamiliar) and regressors for the outcome phase of hit and miss responses for each condition. We also included regressors for error trials during both the delay and outcome phases, as well as six motion parameters for each run and six run constants as regressors of no interest. All the regressors were convolved with a canonical hemodynamic response function. We then applied a high-pass filter (1/128 Hz) to remove low-frequency noise and a first-order autoregressive model to correct for temporal autocorrelation. Beta values for each condition across runs were estimated for all brain voxels and converted to *t* values. We used *t* values in the subsequent analysis, as *t* values have been shown to reduce the influence of noisy voxels and improve performance (Misaki et al., 2010).

### ROI Definition

Based on converging evidence from both animal and human studies demonstrating partner-selectivity in the NAcc (Fujisaki et al., 2026; Ueda & Abe, 2021; Walum & Young, 2018) and the aINS (Fujisaki et al., 2026; Vitale et al., 2025), we defined them as regions of interest (ROIs). Both regions have also been consistently implicated in the anticipation of social rewards, as demonstrated by a meta-analysis of fMRI studies employing the social incentive delay task (Martins et al., 2021). The ROI masks were generated based on the automated anatomical atlas 3 (AAL3) (Rolls et al., 2020) for the NAcc, and the Brainnetome atlas (Fan et al., 2016) for the aINS, given its finer parcellation of insular subregions.

### Multivoxel Pattern Analysis

#### Decoding Analysis

As in our prior studies (Fujisaki et al., 2026; Ueda & Abe, 2021), we conducted pairwise decoding analyses using The Decoding Toolbox (Version 3.999F; Hebart et al., 2015) to examine whether spatial patterns of neural activity in the NAcc and aINS during the anticipation of positive feedback could discriminate the romantic partner from a friend and an unfamiliar individual. For each participant, voxelwise BOLD signal patterns were extracted separately for the left and right hemispheres within each ROI from 18 *t* maps (3 conditions × 6 runs) corresponding to the anticipatory delay phase for each condition and run. Voxelwise *t* values were demeaned across maps to remove condition-nonspecific response components (Op de Beeck, 2010; Walther et al., 2016). Classification between conditions was performed using a linear support vector machine with a cost parameter (C) of 1 and leave-one-run-out cross-validation, and performance was quantified using the area under the curve (AUC). We examined whether classification performance patterns differed by hemisphere using a repeated-measures analysis of variance and found that the interaction between laterality (left and right) and pairing type (partner–friend, partner–unfamiliar, and friend–unfamiliar) was not significant (Table S1); therefore, classification accuracies were averaged across hemispheres within each participant for statistical tests. To test whether the classification performance was significantly above the chance level (AUC = 0.50, reflecting the expected outcome of random predictions in a binary classification task), individual one-sample *t* tests were conducted for each condition pairing, with Bonferroni correction applied to control for multiple comparisons (adjusted *p* = *p* × 3).

#### Multiple Regression RSA

To further examine whether the NAcc and aINS encode romantic partners in a specific manner beyond familiarity, we applied multiple regression RSA (Chikazoe et al., 2014). This approach allowed us to estimate the independent contributions of partner-specific and familiarity-based representational structures to the neural representations (see schematic overview in Figure 2). In this analysis, we first constructed the neural representational dissimilarity matrix (RDM) separately for the left and right hemisphere within each ROI, capturing neural pattern dissimilarity across conditions and runs. This matrix was included into the regression as a dependent variable. Then, we constructed three “model RDMs” representing hypothesized sources of representational structure, which were included into the regression as independent variables. To construct the neural RDM, as in the decoding analyses, we extracted voxelwise activity patterns within each ROI from 18 *t* maps for each participant and demeaned them across maps. We computed pairwise Pearson’s correlations between all condition–run vectors, then converted them to dissimilarity (1 − Pearson’s *r*), resulting in an 18 × 18 RDM. The three model RDMs were binary and defined over the same 18 × 18 set of condition-by-run pairs as the neural RDM. The first one is “Love RDM”, and the second one is “Familiarity RDM”, which were informed by a prior study examining neural representations of personal relevance using the RSA (Bayer et al., 2021). The Love RDM was designed to capture a “partner-specific” representational structure, in which the partner is defined as dissimilar to both the friend and the unfamiliar individual, while the friend and the unfamiliar individual are defined as similar (similar pairs were coded as 0 and dissimilar pairs as 1). The Familiarity RDM was constructed to model familiarity-based representations, defining the partner and the friend as similar and both as dissimilar to the unfamiliar individual. Furthermore, “Session RDM” was included into the regression to account for pattern similarity arising from run-specific shared noise or temporal drift within fMRI sessions (Alink et al., 2015); this RDM coded whether pairs of conditions came from the same run (0) or from different runs (1). We vectorized the lower triangle of each RDM and regressed the neural RDM onto the Love and Familiarity model RDMs while including the Session RDM as a covariate, using ordinary least squares regression. As with the decoding analyses, we found that the interaction between laterality (left and right) and model RDM type (Love RDM and Familiarity RDM) on the regression coefficients was not significant (Table S1); therefore, the *β* coefficients for each RDM were averaged across hemispheres for subsequent analyses. We confirmed that variance inflation factors (VIFs) (Marquardt, 1970) were close to 1 for each of the variables (Love RDM = 1.02, Familiarity RDM = 1.02, Session RDM = 1.04), indicating negligible collinearity among model RDMs, consistent with expectations from standard collinearity diagnostics (Belsley, 1991). Group-level significance of the *β* coefficients for the Love and Familiarity RDMs was assessed using individual one-tailed one-sample *t* tests, evaluating whether the mean *β* coefficient was significantly greater than zero. In this test, family-wise error was controlled using Bonferroni correction across the two *β* coefficients (adjusted *p* = *p* × 2).

**Figure 2.**
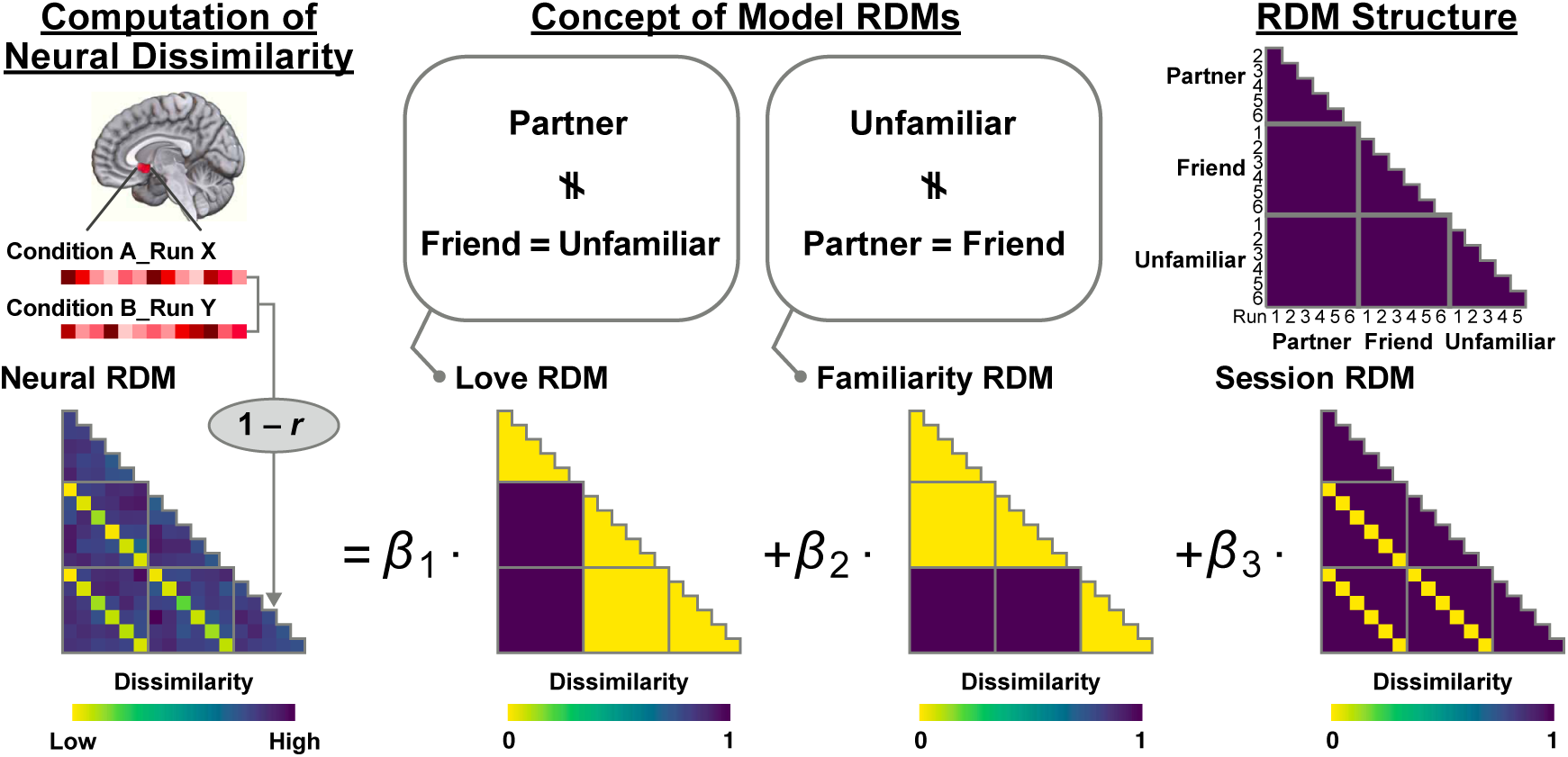
Schematic of multiple regression representational similarity analysis. The neural RDM was constructed by computing pairwise neural dissimilarity (1 – Pearson’s *r*) between extracted neural activity patterns for each condition and run within the region of interest. The model RDMs were defined to reflect hypothesized representational structures, including the Love RDM distinguishing a romantic partner from a friend and an unfamiliar individual, the Familiarity RDM modeling the partner and friend as familiar targets distinct from an unfamiliar individual, and the Session RDM capturing run-related pattern similarity. The regression RSA was performed by modeling the neural RDM as a linear combination of these model RDMs, yielding regression coefficients that quantify the unique contribution of each model to the neural representational structure.

### Correlation Analyses

We further examined associations between relationship duration and partner specificity in the NAcc, and between intrusive thought scores and partner specificity in the aINS. Normality of relationship duration and intrusive thought scores was assessed using the Shapiro–Wilk test. When normality could not be assumed, Spearman’s rank correlation coefficients were used to assess relationships between each behavioral measure and the *β* coefficients for the Love RDM within each ROI. 95% confidence intervals for the correlation coefficients were estimated using a bootstrap procedure with 10,000 resamples, implemented in R using the *boot* package (version 1.3-31; Canty and Ripley, 2025).

Primary correlation analyses were conducted using the full sample; however, when potential outliers were identified, sensitivity analyses excluding these data points were further performed to confirm the robustness of the results. Potential outliers were screened in the *β* coefficients of the Love RDM in the NAcc and aINS, as well as in relationship duration and intrusive thought scores based on two criteria: values exceeding the mean ± 3 SD or those identified by the Smirnov–Grubbs test (Grubbs, 1969).

## Results

### Descriptive statistics of relationship-related measures

Primary descriptive statistics for the relationship-related measures are summarized in Table 1. We confirmed that the subscales for Triangular Love Scale showed good internal consistency (intimacy: Cronbach’s α = 0.80; passion: α = 0.86; commitment: α = 0.79), and that the participants showed relatively high scores on each dimension. In line with the high scores, participants, who were in the early stage of a romantic relationship, generally reported frequent intrusive thoughts (mean = 21.0 %); however, substantial interindividual variability was also observed (SD = 16.6 %).

**Table 1.** Descriptive statistics of relationship-related measures.

| Variable | Mean | SD | Median | Min | Max |
| --- | --- | --- | --- | --- | --- |
| Triangular Love Scale |  |  |  |  |  |
| Intimacy | 7.7 | 0.8 | 7.8 | 5.3 | 9.0 |
| Passion | 6.9 | 1.2 | 7.2 | 4.0 | 8.9 |
| Commitment | 6.7 | 1.3 | 6.7 | 3.3 | 8.7 |
| Number of dating partners | 3.1 | 1.5 | 3 | 1 | 7 |
| Intrusive thought score [%] | 21.0 | 16.6 | 13.3 | 0.0 | 64.1 |

### Partner Advantage across Behavioral Measures

Response times on the social incentive delay task differed significantly across conditions (*F*(2, 100) = 15.66, *P* < 0.001, partial *η*² = 0.24, 95% CI for the effect size = [0.12, 1.00]) (Figure 3A). Participants responded faster in the partner condition than in both the friend condition (*t*(50) = 3.77, adjusted *P* = 0.001, Hedges’ *g* = 0.53, 95% CI for *g* = [0.23, 0.82]) and the unfamiliar condition (*t*(50) = 4.91, adjusted *P* < 0.001, *g* = 0.69, 95% CI = [0.38, 0.99]), which may reflect greater motivation to obtain social approval from their partner. The difference between the friend and unfamiliar conditions did not reach significance (*t*(50) = 2.23, adjusted *P* = 0.091, *g* = 0.31, 95% CI = [0.03, 0.59]).

**Figure 3.**
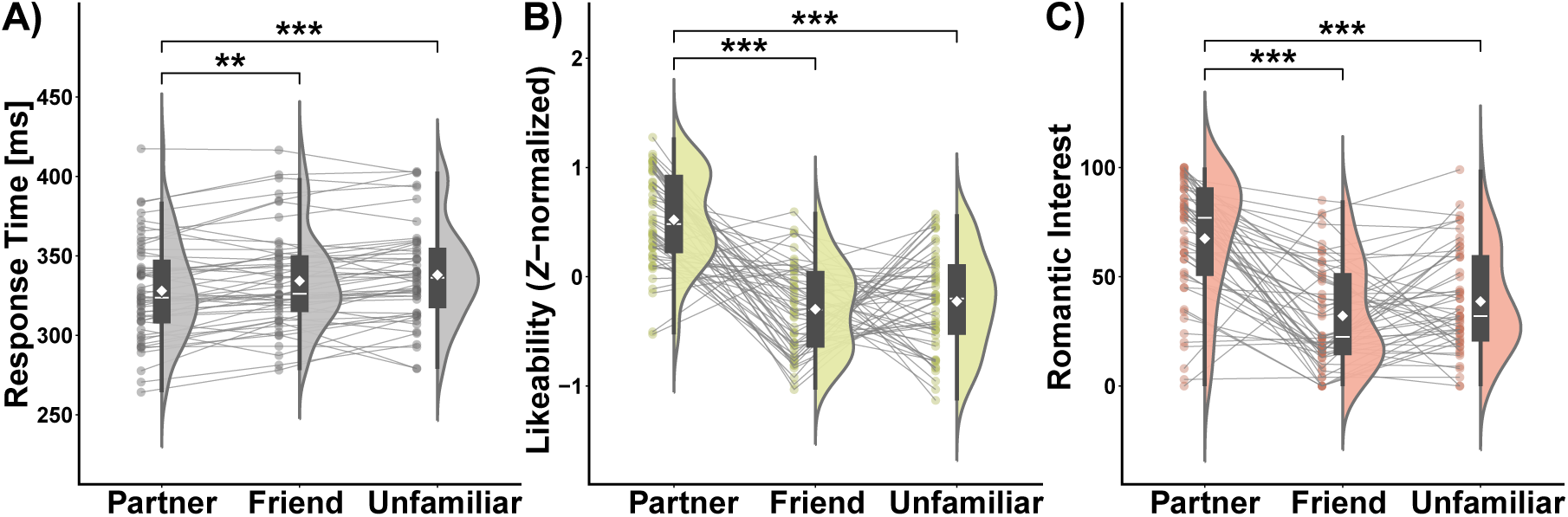
Response times and ratings across conditions. **(A)** Response times were significantly shorter in the partner condition compared with both the friend and unfamiliar conditions. **(B, C)** Participants reported higher likeability ratings for the video clips and greater romantic interest in the partner condition than in the friend and unfamiliar conditions. Across all behavioral panels, half-violin plots depict the data distribution, and boxplots indicate the interquartile range (IQR) with the median shown as a horizontal line. Whiskers extend to values within 1.5 × IQR from the lower and upper quartiles, and white diamond symbols denote mean values. Gray lines link individual participants’ data across conditions (N = 51, except romantic interest: N = 50). \*\**P* < 0.01, \*\*\**P* < 0.001, Bonferroni-corrected.

Consistent with these results, likeability ratings toward the video clips also differed by condition (*F*(2, 100) = 37.83, *P* < 0.001, partial *η*² = 0.43, 95% CI = [0.31, 1.00]) (Figure 3B). Participants rated their partner as more likeable than both the friend (*t*(50) = 8.23, adjusted *P* < 0.001, *g* = 1.15, 95% CI = [0.79, 1.50]) and the unfamiliar individual (*t*(50) = 6.85, adjusted *P* < 0.001, *g* = 0.96, 95% CI = [0.62, 1.29]). Ratings for the friend versus the unfamiliar individual did not differ significantly (*t*(50) = 0.69, adjusted *P* = 1.00, *g* = 0.10, 95% CI = [0.18, 0.37]). Romantic interest ratings yielded a similar pattern (*F*(2, 98) = 35.21, *P* < 0.001, partial *η*² = 0.42, 95% CI = [0.29, 1.00]) (Figure 3C). Participants reported greater romantic interest in their partner than in the friend (*t*(49) = 7.43, adjusted *P* < 0.001, *g* = 1.05, 95% CI = [0.70, 1.39]) and the unfamiliar individual (*t*(49) = 6.03, adjusted *P* < 0.001, *g* = 0.85, 95% CI = [0.53, 1.17]). The difference between the friend and the unfamiliar individual was not significant (*t*(49) = 1.70, adjusted *P* = 0.28, *g* = 0.24, 95% CI = [−0.04, 0.52]). Across all measures, participants showed a consistent partner advantage in both response times and subjective ratings, indicating a clear preference for the partner accompanied by heightened motivation to obtain positive social feedback. These results also align with previous studies (Fujisaki et al., 2026; Ueda & Abe, 2021).

### Neural Differentiation of Partners from Friends and Unfamiliar Individuals in the NAcc and aINS

As in our previous work (Fujisaki et al., 2026; Ueda & Abe, 2021), we first examined whether pairwise decoding of neural activity patterns could discriminate the romantic partner from a close friend and an unfamiliar individual in the NAcc and aINS. In the NAcc, classification accuracy was significantly above chance for both the partner–friend pairing (mean AUC = 0.55, SD = 0.15, *t*(50) = 2.54, adjusted *P* = 0.021, *g* = 0.36, 95% CI for *g* = [0.07, 0.64]) and the partner–unfamiliar pairing (mean = 0.56, SD = 0.16, *t*(50) = 2.67, adjusted *P* = 0.015, *g* = 0.37, 95% CI = [0.09, 0.66]), whereas the friend–unfamiliar pairing did not reach significance (mean = 0.52, SD = 0.13, *t*(50) = 1.11, adjusted *P* = 0.41, *g* = 0.15, 95% CI = [–0.12, 0.43]) (Figure 4A). In the aINS, classification accuracy was significantly above chance for both the partner–friend (mean = 0.62, SD = 0.18, *t*(50) = 4.62, adjusted *P* < 0.001, *g* = 0.65, 95% CI = [0.34, 0.95]) and partner–unfamiliar pairings (mean = 0.62, SD = 0.19, *t*(50) = 4.47, adjusted *P* < 0.001, *g* = 0.63, 95% CI = [0.32, 0.92]) (Figure 4B). In contrast to the NAcc, decoding accuracy for the friend–unfamiliar pairing in the aINS was also above chance (mean = 0.54, SD = 0.14, *t*(50) = 2.26, adjusted *P* = 0.042, *g* = 0.32, 95% CI = [0.03, 0.60]). These results indicate that partner representations in both ROIs could be reliably distinguished from those of other individuals, consistent with our previous findings (Fujisaki et al., 2026; Ueda & Abe, 2021).

**Figure 4.**
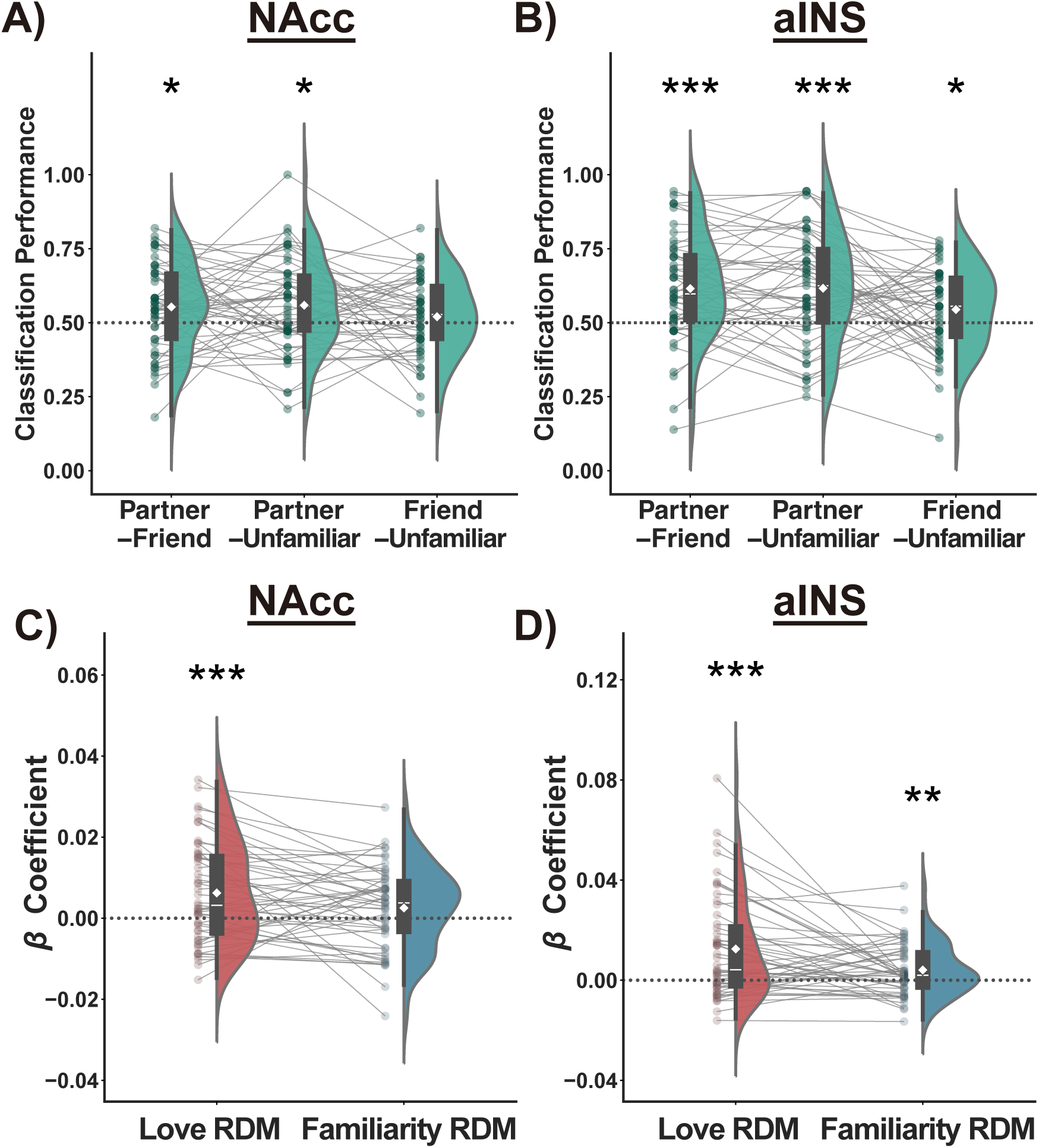
Results of decoding analysis and multiple regression RSA in the NAcc and aINS. **(A, B)** Classification performance for both the partner–friend pairing and the partner–unfamiliar pairing was significantly above chance level (indicated by the dotted line) in the NAcc and aINS. **(C, D)** The *β* coefficients for the Love RDM were significantly greater than zero in both regions. Across all panels, half-violin plots depict the data distribution, and boxplots indicate the interquartile range (IQR) with the median shown as a horizontal line. Whiskers extend to values within 1.5 × IQR from the lower and upper quartiles, and white diamond symbols denote mean values. Gray lines link individual participants’ data across conditions (N = 51). \**P* < 0.05, \*\**P* < 0.01, \*\*\**P* < 0.001, Bonferroni-corrected.

While our primary analyses focused on the NAcc and aINS, we explored dorsal striatal regions, including the caudate nucleus and putamen, where prior work has reported distinguishable neural representations of romantic partners and friends (Fujisaki et al., 2026). Because these analyses were exploratory and not specified in our primary hypotheses, the corresponding results are reported in the Supplementary Material. Briefly, these analyses replicated prior findings by showing that neural representations of the romantic partner were discriminable from those of others in the dorsal striatal regions.

### Partner-Specific Representations Beyond Familiarity in the NAcc and aINS

We next conducted a multiple regression RSA to estimate the unique contribution of partner specificity to neural representations in the NAcc and aINS. This approach allows for an examination of partner specificity in each region beyond familiarity. In the NAcc, the *β* coefficient for the Love RDM was significantly positive (*t*(50) = 3.57, adjusted *P* < 0.001, *g* = 0.50, 95% CI for *g* = [0.21, 0.79]), whereas the *β* coefficient for the Familiarity RDM was not (*t*(50) = 1.76, adjusted *P* = 0.084, *g* = 0.25, 95% CI = [–0.03, 0.52]) (Figure 4C). In the aINS, the *β* coefficients for both the Love and the Familiarity RDMs were significantly positive (Love: *t*(50) = 4.28, adjusted *P* < 0.001, *g* = 0.60, 95% CI = [0.30, 0.90]; Familiarity: *t*(50) = 2.77, adjusted *P* = 0.008, *g* = 0.36, 95% CI = [0.10, 0.67]) (Figure 4D). These results indicate that partner-specific representational structure is present in both ROIs, with familiarity-related structure also represented in the aINS.

Building on the exploratory results from the decoding analysis showing distinguishable representations of the partner in the caudate nucleus and putamen, we conducted multiple regression RSA in these regions. We found that partner-specific representational structure was also observed in both the caudate nucleus and putamen, with familiarity-related representations additionally observed in the putamen (see Supplementary Material for detailed results).

### Attenuated Partner Specificity in the NAcc with Longer Relationship Duration

Building on our prior work showing that the distinctiveness of partner representations in the NAcc is negatively associated with relationship duration (Fujisaki et al., 2026), we examined whether this association would also be observed even after controlling for familiarity-based representations. To this end, we assessed whether relationship duration was related to the *β* coefficient for the Love RDM in the NAcc. Given that relationship duration showed a significant departure from normality (Shapiro–Wilk test, *P* < 0.001), we used Spearman’s rank correlation for this analysis. This analysis revealed that relationship duration was negatively associated with the Love RDM *β* (*ρ* = −0.38, *P* = 0.006, 95% CI = [–0.58, –0.12]) (Figure 5A), indicating that consistent with our previous findings (Fujisaki et al., 2026), individuals with longer relationship durations tend to show reduced representational specificity for the romantic partner in the NAcc.

**Figure 5.**
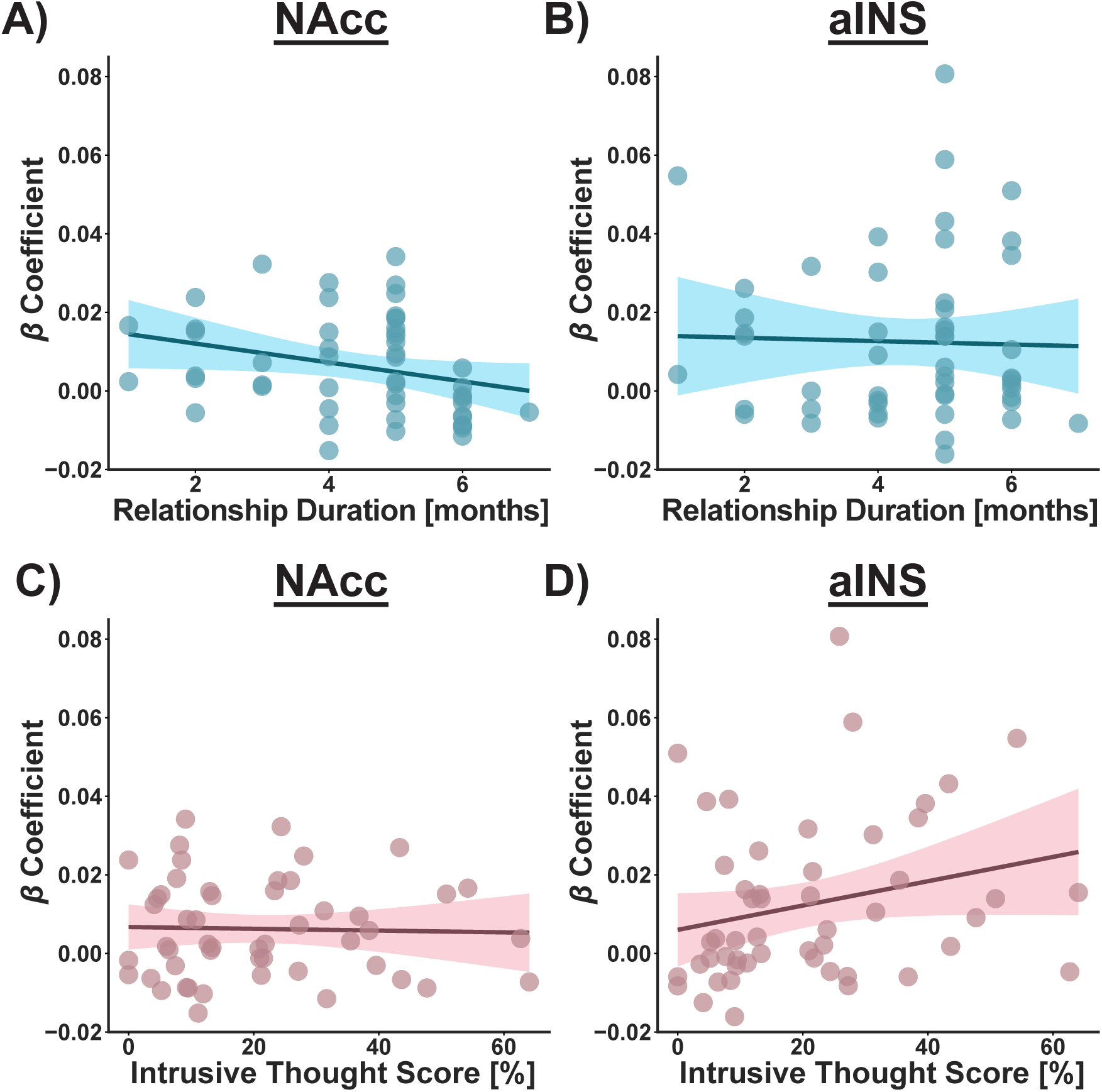
Results of correlation analyses in the NAcc and aINS. **(A, B)** Relationship duration is negatively associated with the *β* coefficient for the Love RDM in the NAcc but not in the aINS. **(C, D)** Intrusive thought scores were positively associated with the *β* coefficient for the Love RDM in the aINS but not in the NAcc. Solid lines represent the fitted regression lines, with shaded areas illustrating the 95% confidence intervals (N = 51).

To assess the regional specificity of this association, we conducted exploratory analyses in the aINS that also represented the partner in a specific way beyond familiarity. Consistent with prior work (Fujisaki et al., 2026), no comparable associations with relationship duration were observed in the aINS (*ρ* = −0.03, *P* = 0.84, 95% CI = [–0.30, 0.26]) (Figure 5B). We further directly compared the strength of the association between the NAcc and the aINS using one-tailed Steiger’s tests (Steiger, 1980), which revealed a significantly stronger association in the NAcc than in the aINS (*P* = 0.032), indicating that the attenuation of partner-specific representations with increasing relationship length was specific to the NAcc. This regional specificity was further corroborated by our exploratory analyses in the caudate nucleus and putamen, which showed non-significant correlations with relationship duration that were significantly weaker than those in the NAcc (see Supplementary Material).

### Greater Partner Specificity in the aINS Relates to More Frequent Intrusive Thoughts about the Partner

We next examined whether individuals who exhibited greater partner specificity in the aINS more frequently experienced intrusive thoughts about their partner. As intrusive thought scores deviated from normality (Shapiro–Wilk test, *P* < 0.001), we again employed Spearman’s rank correlation. Consistent with our hypothesis, the *β* coefficient for the Love RDM in the aINS showed a significant positive association with intrusive thought scores (*ρ* = 0.30, *P* = 0.031, 95% CI = [0.002, 0.57]; Figure 5D). We also confirmed that this association remained significant even after excluding one participant identified as an outlier in the *β* estimate (see Supplementary Material). These results indicate that more distinctive partner-specific representations in the aINS were associated with more frequent intrusive thoughts about the partner. In contrast to the aINS, frequency of intrusive thoughts was not significantly associated with partner specificity in the NAcc (*ρ* = 0.02, *P* = 0.90, 95% CI = [–0.26, 0.31]) (Figure 5C), and the association with intrusive thought scores tended to be stronger in the aINS than in the NAcc (one-tailed Steiger’s test, *P* = 0.069). The regional specificity of the aINS was further substantiated by exploratory analyses in the caudate nucleus and putamen, which showed weaker and non-significant associations with intrusive thought scores compared to those observed in the aINS (see Supplementary Material).

## Discussion

The present study investigated whether neural representations in the NAcc and aINS encode a partner in a manner dissociable from a friend and an unfamiliar attractive individual. We employed multiple regression RSA, which allows for testing the unique contribution of partner specificity beyond familiarity to neural representations, and found that both the NAcc and aINS exhibit partner-specific representations. Our investigation first contributes by replicating previous findings (Fujisaki et al., 2026; Ueda & Abe, 2021) with improved methodology that addresses prior limitations regarding potential effects derived from familiarity. Moreover, consistent with our previous study (Fujisaki et al., 2026), we observed that the partner specificity in the NAcc was attenuated among participants with longer relationship duration. This effect was not observed in other regions that encoded the partner in a specific way. As a novel finding, we observed that participants who exhibited greater partner specificity in the aINS experienced more frequent intrusive thoughts about the partner. Together, these findings suggest that while both the NAcc and aINS engage in partner-specific processing, the psychological and functional aspects supported by such processing may differ across these regions.

The primary finding of the present study is that partner-specific neural representations in the NAcc and aINS are not reducible to differences in familiarity with others. This issue was not directly tested in our previous studies using the same experimental procedures (Fujisaki et al., 2026; Ueda & Abe, 2021). By explicitly modeling both partner specificity and familiarity in a multiple regression RSA framework, we revealed that the partner is represented in the NAcc and aINS in a manner dissociable from a friend and an unfamiliar attractive individual. This finding further implies that romantic love is represented not along a single dimension of social closeness or familiarity, but as a qualitatively distinct form of social relationship. Enhanced NAcc and aINS responses to familiar and socially relevant targets, including friends and family members, have been widely reported (e.g., Bortolini et al., 2024); however, from the perspective of representational similarity, romantic partners appear to be represented differently from a wide range of other close interpersonal relationships. This account is also consistent with evidence from prairie voles showing that experience-dependent plasticity in the NAcc and aINS is observed in individuals that have formed a pair bond, but not in individuals cohabiting with same-sex siblings despite comparable familiarity (Aragona et al., 2003, 2006; Vitale et al., 2025). Together, these findings suggest that the human capacity for forming and maintaining romantic relationships may be supported by neural mechanisms that encode partners as qualitatively distinct social targets.

We further observed that the distinctiveness of partner representations in the NAcc was negatively associated with relationship length. This result replicates our previous study (Fujisaki et al., 2026) under a modified experimental design that explicitly controlled for familiarity. Considering that participants in the current study were in the early stage of a romantic relationship (within six months of relationship initiation), the present findings suggest that partner-specific representations in the NAcc are established early in romantic relationships but are modulated within this period. The observed decrease in partner specificity in the NAcc likely reflects a reduction in the novelty or reward value of partner-related stimuli as the relationship progresses. This interpretation aligns with the well-established role of the NAcc in assigning incentive value to novel and highly rewarding stimuli (Berridge & Robinson, 1998; Salamone & Correa, 2012). A persistent attraction to the partner has been thought to be one of the key components for partner preference and relationship maintenance (Feldman, 2017; Walum & Young, 2018). Notably, consistent with our prior work (Fujisaki et al., 2026), exploratory analyses revealed that such a negative association with relationship length was not observed in the dorsal striatal regions. Given the established role of the dorsal striatum in the learning and maintenance of habitual actions (Balleine et al., 2007; Balleine & O’Doherty, 2010), this regional dissociation suggests that partner representations in the NAcc are particularly sensitive to changes in the novelty or reward value of partner-related stimuli over time, whereas representations in dorsal striatal regions may be relatively stable across relationship progression, at least within six months of relationship initiation. This possibility is consistent with theoretical predictions proposing that functional reliance may shift from the ventral to dorsal striatum, reflecting a transition from reward- and novelty-driven processing to habit-like processes that support the persistence of partner preferences and behaviors (Burkett et al., 2011; Feldman, 2017; Tops et al., 2014). However, this interpretation remains speculative, and further research will be required to directly test these proposed mechanisms.

We found that greater partner specificity in the aINS was positively associated with more frequent intrusive thoughts about the partner. This finding provides the first empirical evidence linking partner representations in the aINS to the intrusive and obsessive cognitive aspects of early romantic love. Notably, data on intrusive thoughts were quantitatively derived from participants’ actual everyday experiences in real-life contexts, rather than from retrospective self-reports used in previous studies (Langeslag et al., 2012; Nilakantan et al., 2014). Given that the aINS is necessary for conscious urges and cravings in substance addiction (Naqvi et al., 2007; Naqvi & Bechara, 2009) and that it is thought to recall and maintain salient representations in mind (Naqvi et al., 2014; Naqvi & Bechara, 2010), our results suggest that the aINS may perform an analogous function in romantic relationships. Specifically, partner-related information may be represented with heightened salience in the aINS, contributing to the frequent intrusion of partner-related thoughts. This possibility is consistent with converging evidence implicating the aINS in OCD (de Wit et al., 2014; Norman et al., 2019; Yu et al., 2022). Romantic love and OCD share a core phenomenological feature—intrusive and repetitive thoughts focused on a specific concern (Doron et al., 2014; Fisher et al., 2016; Leckman et al., 1999; Thompson et al., 2020)—and have been reported to exhibit partially overlapping neurochemical profiles (Marazziti et al., 1999, 2015; Schneiderman et al., 2012). Together, these observations suggest that partially overlapping neural processes may contribute to intrusive and obsessive thoughts in romantic love and OCD. In this light, the present findings indicate that the involvement of the aINS, previously implicated in the context of OCD, may extend beyond pathological states to support intrusive partner-related thoughts in romantic relationships.

This study has two limitations that suggest important directions for future research. First, our sample was restricted to heterosexual men to avoid potential sex-related confounds and to maintain methodological consistency with previous research (Fujisaki et al., 2026; Ueda & Abe, 2021); however, the generalizability across sex and sexual orientation is limited. Second, the cross-sectional design precludes conclusions about how partner-specific representations develop over time. To fully capture the temporal dynamics of romantic bonding, future studies should adopt longitudinal designs that follow more diverse sex and sexual orientations across different stages of their relationships.

In conclusion, the present study provides evidence that partner-specific neural representations in the NAcc and aINS relate to distinct aspects of romantic bonding. Using multiple regression RSA, we demonstrate that the NAcc and aINS encode partner specificity beyond familiarity, with partner specificity in the NAcc established early and modulated by relationship duration, and in the aINS associated with intrusive thoughts about the partner. These findings demonstrate that partner-specific neural representations in the NAcc and aINS reflect dissociable functions, suggesting that romantic bonding engages distinct neural processes that may together support the selective commitment toward a romantic partner.

## Data and Code Accessibility

The data and code are available at: https://doi.org/10.5281/zenodo.18628958

## Acknowledgments

We thank Maki Terao and Aiko Murai for their technical support in MRI data acquisition. This study was conducted using the MRI scanner and related facilities at the Institute for the Future of Human Society, Kyoto University.

