## Supplementary Material for "How the brain represents a romantic partner: dissociable roles of the nucleus accumbens and anterior insula"

### 1. Exploratory analyses in the caudate nucleus and putamen

Our prior work using decoding analyses demonstrated that the caudate nucleus and putamen showed distinguishable representations of partners from friends (Fujisaki et al., 2026). Given these findings, we conducted exploratory analyses to investigate whether the dorsal striatal regions encode romantic partners in a specific manner beyond familiarity. The masks for the caudate nucleus and putamen were generated using the automated anatomical labeling atlas 3 (Rolls et al., 2020). As in the NAcc and aINS, no discernible hemispheric differences were observed in classification performance or in the $\beta$ coefficients for the Love RDM, and measures were therefore averaged across hemispheres for subsequent analyses (**Table S1**).

Decoding analyses replicated previous findings (Fujisaki et al., 2026) by demonstrating that neural representations of the romantic partner could be distinguished from those of other individuals (**Figure S1A, B**). Specifically, classification accuracy in both the caudate nucleus and putamen exceeded chance level for the partner–friend pairing (caudate: mean AUC = 0.57, SD = 0.16, $t(50) = 3.08$, adjusted $P = 0.005$, $g = 0.43$, 95% CI for $g$ = [0.14, 0.72]; putamen: mean = 0.58, SD = 0.14, $t(50) = 4.18$, adjusted $P < 0.001$, $g = 0.59$, 95% CI = [0.29, 0.88]) and the partner–unfamiliar pairing (caudate: mean = 0.60, SD = 0.17, $t(50) = 2.54$, adjusted $P = 0.021$, $g = 0.36$, 95% CI = [0.30, 0.89]; putamen: mean = 0.62, SD = 0.14, $t(50) = 5.78$, adjusted $P < 0.001$, $g = 0.81$, 95% CI = [0.49, 1.12]), whereas classification performance did not exceed chance for the friend–unfamiliar pairing in either region (caudate: mean = 0.51, SD = 0.17, $t(50) = 0.62$, adjusted $P = 0.81$, $g = 0.09$, 95% CI = [–0.19, 0.36]; putamen: mean = 0.53, SD = 0.16, $t(50) = 1.51$, adjusted $P = 0.21$, $g = 0.21$, 95% CI = [–0.07, 0.49]).

Multiple-regression RSA further revealed that the caudate nucleus and putamen encode partner-specific representations beyond familiarity (Figure S1C, D). In the caudate nucleus, we observed a significant positive $\beta$ coefficient for the Love RDM ($t(50) = 3.13$, adjusted $P = 0.003$, $g = 0.44$, 95% CI for $g$ = [0.15, 0.72]), while the $\beta$ coefficient for the Familiarity RDM did not reach significance ($t(50) = 1.76$, adjusted $P = 0.084$, $g = 0.25$, 95% CI = [–0.03, 0.52]) (**Figure S1C**). In the putamen, $\beta$ estimates were significantly greater than zero for both the Love RDM ($t(50) = 4.93$, adjusted $P < 0.001$, $g = 0.69$, 95% CI = [0.38, 0.99]) and the Familiarity RDM ($t(50) = 3.54$, adjusted $P <$ 0.001, $g = 0.50$, 95% CI = [0.20, 0.78]) (**Figure S1D**). These results indicate that both the caudate nucleus and the putamen encode partner-specific representational structure beyond familiarity, broadly paralleling the patterns observed in the NAcc and aINS.

We also conducted exploratory correlation analyses between the $\beta$ coefficients for the Love RDM in the dorsal striatal regions and both relationship duration and intrusive thought scores, to examine whether the associations observed in the NAcc and aINS were specific to these regions and not present in the caudate nucleus and putamen. Importantly, neither the caudate nucleus nor the putamen showed significant correlations with relationship duration (caudate: $\rho = -0.11$, $P = 0.43$, 95% CI = [–0.37, 0.16]; putamen: $\rho = -0.04$, $P = 0.78$, 95% CI = [–0.34, 0.26]; **Figure S2A, B**) or with intrusive thought scores (caudate: $\rho = -0.002$, $P = 0.99$, 95% CI = [–0.28, 0.28]; putamen: $\rho = 0.02$, $P = 0.89$, 95% CI = [–0.24, 0.28]; **Figure S2C, D**). We also examined whether the strength of the observed correlations in the NAcc and aINS was greater than that in the caudate nucleus and putamen using one-tailed Steiger's tests (Steiger, 1980). These analyses showed that the association with relationship duration was significantly stronger in the NAcc than in both the caudate nucleus ($P = 0.047$) and putamen ($P = 0.021$). Similarly, the association with intrusive thoughts was significantly stronger in the aINS than in the caudate nucleus ($p = 0.007$) and the putamen ($P = 0.030$). Taken together, although partner-specific representational structure was observed in the dorsal striatal regions, the absence of associations with relationship duration and intrusive thoughts indicates that these regions are functionally dissociated from the NAcc and aINS.

### 2. Correlation analyses in the aINS after excluding an outlier

To assess whether the observed correlations in the NAcc and aINS were driven by outliers, we conducted robustness analyses. For the $\beta$ coefficients of the Love RDM in the NAcc and aINS, as well as relationship duration and intrusive thought scores, potential outliers were identified using two criteria: values exceeding the mean ± 3 standard deviations and those detected by the Smirnov–Grubbs test. One participant met both outlier criteria for the Love RDM $\beta$ in the aINS ($\beta = 0.081$, exceeding the mean + 3 standard deviations threshold of 0.075; Smirnov–Grubbs test, $P = 0.027$). No participants met the outlier criteria for any of the other variables. We then repeated the correlation analysis between intrusive thoughts and the $\beta$ coefficients for Love RDM in the aINS after excluding this participant, and found that the association remained statistically significant ($\rho = 0.29$, $P = 0.040$, 95% CI = [–0.02, 0.57]). These results indicate that the association between aINS partner-specificity and intrusive thoughts remained robust after excluding the identified outlier.

**Table S1. Results of repeated-measures analyses of variance testing interactions between laterality and pairing type for classification accuracy and between laterality and model RDM type for $\beta$ coefficients in each region.**

| Variable | Region | *F* | Df | *P* value |
|---|---|---|---|---|
| Classification Accuracy | NAcc | 0.59 | (2.0, 98.3) | 0.56 |
| | aINS | 0.54 | (1.7, 85.5) | 0.56 |
| | Caudate | 1.15 | (1.9, 96.2) | 0.32 |
| | Putamen | 0.64 | (1.9, 97.1) | 0.53 |
| Regression Coefficient | NAcc | 0.98 | (1, 50) | 0.33 |
| | aINS | 0.07 | (1, 50) | 0.79 |
| | Caudate | 2.57 | (1, 50) | 0.12 |
| | Putamen | 0.70 | (1, 50) | 0.41 |

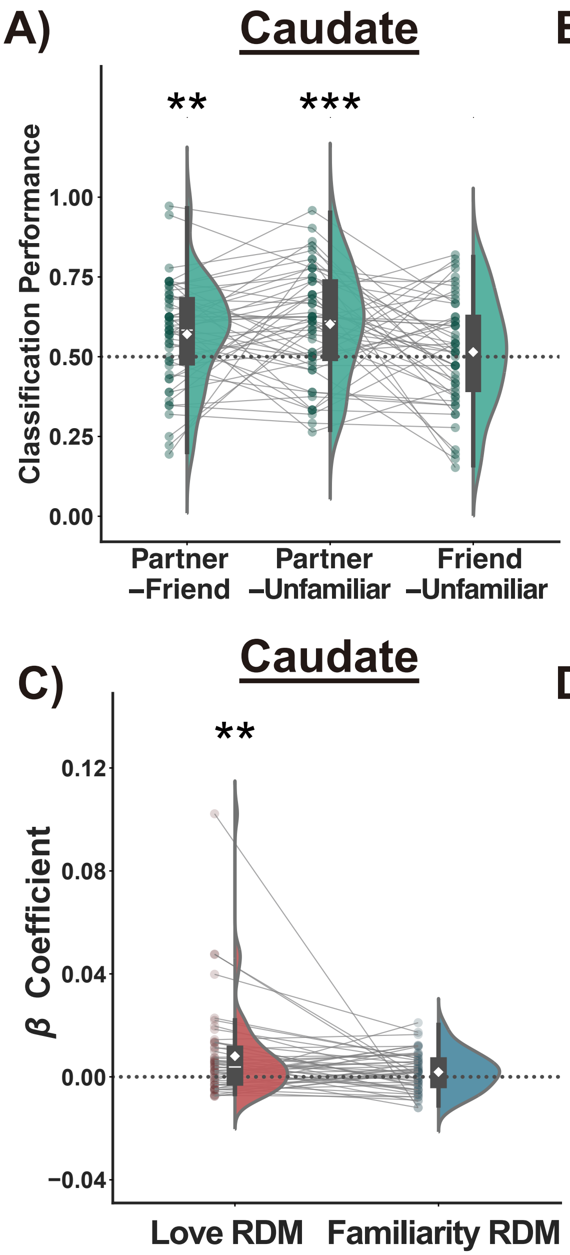


**Figure S1. Results of decoding analysis and multiple regression RSA in the caudate nucleus and putamen.**

**(A, B)** Classification performance for both the partner–friend pairing and the partner–unfamiliar pairing was significantly above chance level (indicated by the dotted line) in the caudate nucleus and putamen.

**(C, D)** The $\beta$ coefficients for the Love RDM were significantly greater than zero in both regions. Across all panels, half-violin plots depict the data distribution, and boxplots indicate the interquartile range (IQR) with the median shown as a horizontal line. Whiskers extend to values within 1.5 × IQR from the lower and upper quartiles, and white diamond symbols denote mean values. Gray lines link individual participants' data across conditions (N = 51). *$P < 0.05$, **$P < 0.01$, ***$P < 0.001$, Bonferroni-corrected.

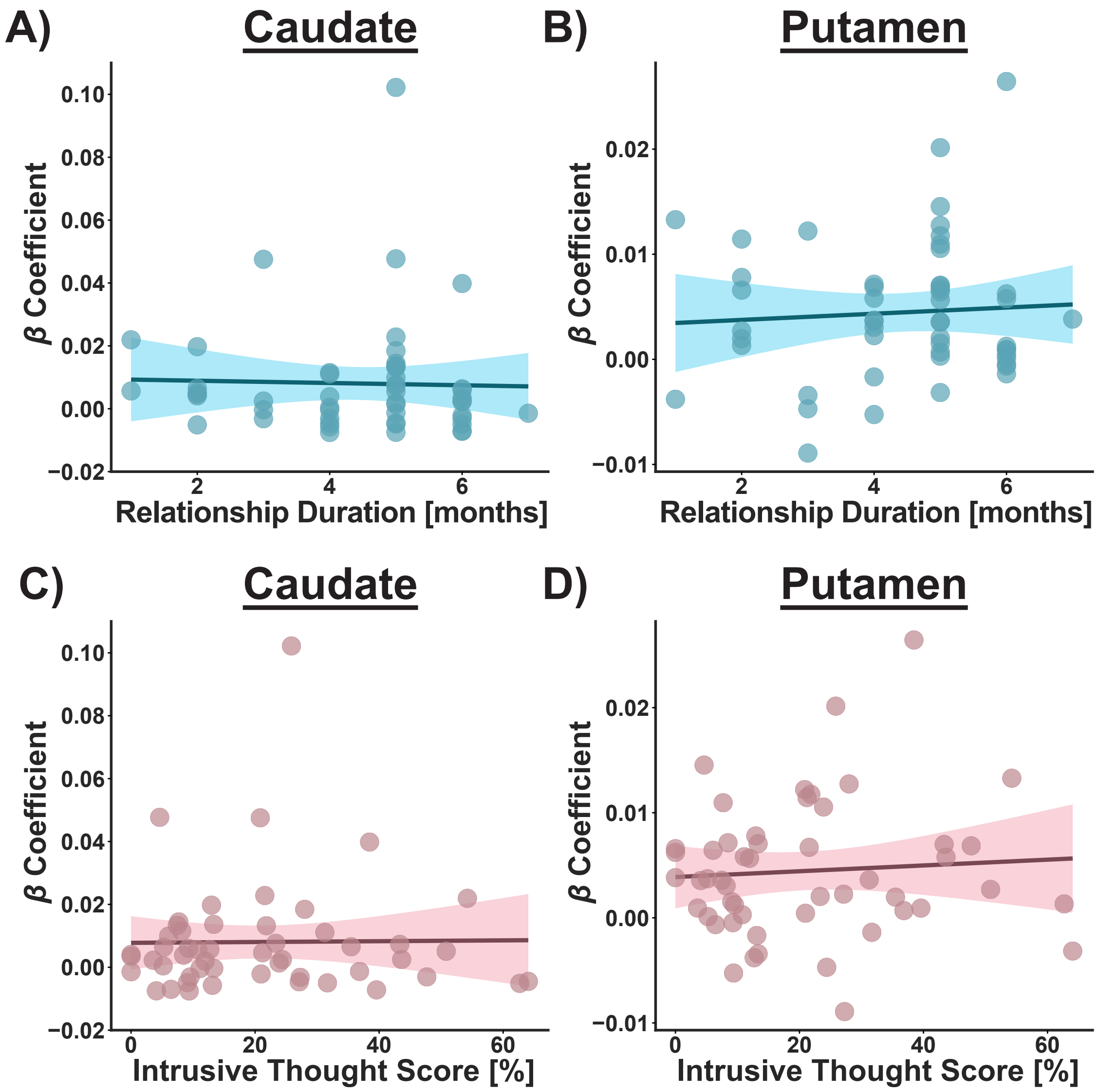


**Figure S2. Results of correlation analyses in the caudate nucleus and putamen.** **(A–D)** Neither relationship duration nor intrusive thought scores showed significant associations with the $\beta$ coefficients for the Love RDM in the caudate nucleus or putamen. Solid lines represent the fitted regression lines, with shaded areas illustrating the 95% confidence intervals (N = 51).
